# Perturbing human V1 degrades the fidelity of visual working memory

**DOI:** 10.1101/2024.06.19.599798

**Authors:** Mrugank Dake, Clayton E. Curtis

## Abstract

Decades of macaque research established the importance of prefrontal cortex for working memory. Surprisingly, recent human neuroimaging studies demonstrated that the contents of working memory can be decoded from primary visual cortex (V1). However the necessity of this mnemonic information remains unknown and contentious. Here we provide causal evidence that transcranial magnetic stimulation targeting human V1 disrupted the fidelity of visual working memory. Errors increased only for targets remembered in the portion of the visual field disrupted by stimulation. Moreover, concurrently measured electroencephalography confirmed that stimulation disrupted not only memory behavior, but neurophysiological signatures of working memory. These results change the question from whether visual cortex is necessary for working memory to what mechanisms it uses to support memory. Moreover, they point to models in which the mechanisms supporting working memory are distributed across brain regions, including sensory areas that here we show are critical for memory storage.

## Introduction

Our ability to maintain information no longer present in the environment, referred to as working memory, acts as a building block supporting most higher forms of cognition ^1^. Indeed, impaired working memory can cause a cascade of broad cognitive symptoms in psychiatric disease ^2^. Based on pioneering studies, the prefrontal cortex is theorized to store information in working memory in recurrent networks whose neural activity forms a mnemonic bridge between perception and memory-guided behavior ^3,4^. There are two major pillars of support for this theory from studies of macaques. One, activity in prefrontal cortical neurons persists during memory delays ^5^. Two, lesions to the macaque prefrontal cortex impair working memory performance ^6^. More recently, human neuroimaging studies demonstrate that working memory representations may be widely distributed in the brain beyond the prefrontal cortex ^7,8^, including evidence for persistent neural activity in many cortical ^9^ and even subcortical areas ^10^. Similarly, lesions to human cortical areas, both within and outside of the prefrontal cortex, impact working memory performance ^11,12^.

The most surprising finding in working memory research, which has now been replicated numerous times, is that the multivariate patterns of neural activity in primary visual cortex, V1, can be used to decode the contents of visual working memory ^13,14^. Moreover, patterns in V1 are often the best for decoding the contents of working memory. Therefore, activity in V1 may encode representations needed for storing visual features in working memory ^15–17^, an idea that has been termed the sensory recruitment model of working memory. However, healthy skepticism exists about the necessity of V1 for working memory, as both macaque neurophysiology and human neuroimaging are mixed as to whether the neural activity in V1 persists during retention intervals ^9,18^, a hallmark of working memory. Perhaps the information decoded from V1 is not necessary for working memory despite being present. Lesions cannot resolve whether V1 is necessary for working memory because lesions to V1 would simply prevent visual encoding.

To overcome this impasse, here we directly tested whether human V1 is necessary for working memory using transcranial magnetic stimulation (TMS) to perturb neural processing during retention intervals.

## Results

We first used fMRI to identify and model the receptive field properties of V1 ^19^. Then, we identified the portion of V1 accessible to target with TMS - the dorsal part of the retinotopic map that represents the contralateral lower quadrant of the visual field (Figure 1A). We modeled the electric field induced by TMS ^20^ to target V1 in each participant (Figure 1B; see individual E-field maps in Supp Figure 1). These models indicated that TMS induced a greater electrical field in the V1 hemisphere targeted (Figure 1C). Although we directed the focus of the TMS pulse to V1, our electric field models suggested that to a lesser extent dorsal V2 may have also been affected. Targeting V1 with TMS induced phosphenes whose spatial extents were drawn by participants. This allowed us to identify the phosphene field and its mirrored location (Figure 1D). Phosphene fields aligned well with the biophysical models we used to estimate the electric fields induced by TMS.

**Figure 1.**
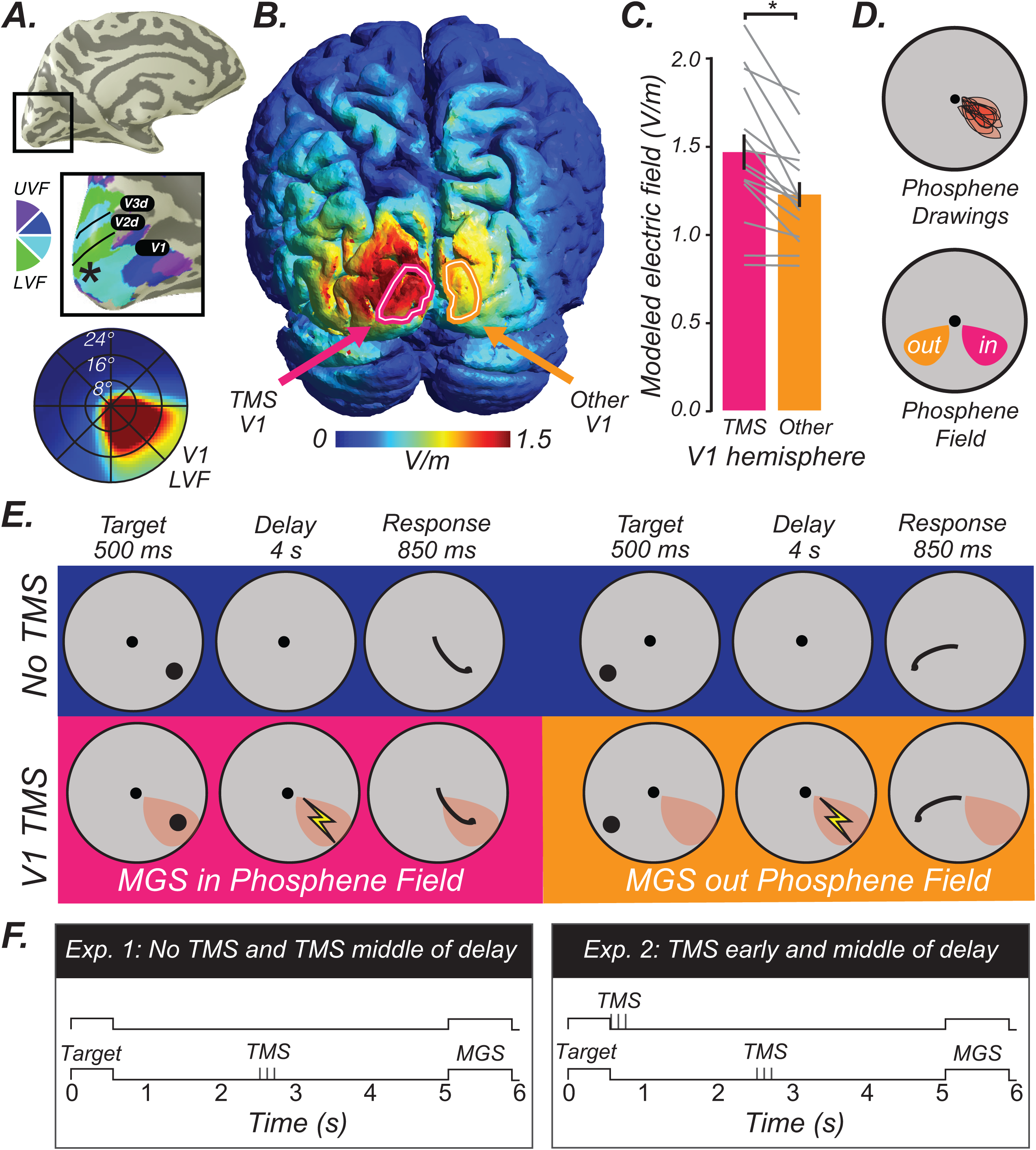
Working memory task conditions, trial types, and TMS procedures. **A.** In each participant, V1 was identified using population receptive field mapping ^19^. Dorsal V1, which represents the contralateral lower visual field (LVF), was accessible to TMS stimulation (asterisk marks TMS target), while the upper visual field (UVF) is buried deeper and not accessible. The bottom radial plot depicts the group average visual field coverage of the dorsal V1 accessible to TMS, whose pRFs are localized to the lower visual field. **B.** The modeled electrical field induced by TMS targeting the retinotopically-defined left V1 (magenta encircled) for an example participant. V1 in the hemisphere not perturbed with TMS encircled by an orange line. See Supp Figure 1 for simulated electrical fields for all participants. Simulated electric field in Volts/meter (V/m). **C.** Based on the model ^20^, TMS induced a significantly larger electric field in the stimulated V1 compared to the V1 in the other hemisphere, (paired-*t* test, two-sided, *t*(14)=4.677, *p=0.00035, d=2.50, n=15* humans). Thin gray lines are individual participants (n=15), bars are means (+/- SEM) across participants. **D.** In each participant, phosphene fields were defined as the portion of the visual field in which phosphenes were reliably evoked. TMS to dorsal V1 produced phosphenes in the contralateral lower visual field. Prior to memory experiments, participants drew with a mouse the extent of TMS-induced phosphenes (top). During the memory experiments, we placed visual targets in the phosphene field, and out of it (in the mirrored location) (bottom). **E.** Participants performed a memory-guided saccade (MGS) task, during which they maintained in working memory a target location. After a retention interval of 4 seconds, participants generated a memory-guided saccade. Feedback was then given, followed by an intertrial interval (2-3 s; not shown). We measured the accuracy of memory-guided saccades for targets in the phosphene field (experimental condition; magenta). Control conditions included no TMS (blue) and TMS to V1 when targets were in the hemifield opposite to the phosphene field (orange). **F**. Trial schematics for Experiment 1 and Experiment 2. Data are provided in the Source Data file.

In Experiment 1, we measured the accuracy of memory-guided saccades for targets placed within the phosphene field (i.e., contralesional to TMS), and compared the performance to that when no TMS was applied and to that when the targets were placed in the hemifield opposite the phosphene field (i.e., ipsilesional to TMS) (Figure 1E/F). Remarkably, TMS applied to V1 during the middle of the delay caused larger memory errors, but only for targets within the phosphene field (Figure 2A). Specifically, memory errors were greater for targets inside the phosphene field compared to when no TMS was applied (*t*(14)=2.431, *p*=0.013, *d*=1.30), and compared to targets in the mirrored hemifield of the phosphene field (*t*(14)=2.506, *p*=0.012, *d*=1.34). Memory errors were not different for targets outside the phosphene field during TMS and during no TMS trials (*t*(14)=-0.678, *p*=0.745, *d*=0.36) confirming the retinotopic specificity of the effect of TMS targeting V1 on working memory performance.

**Figure 2.**
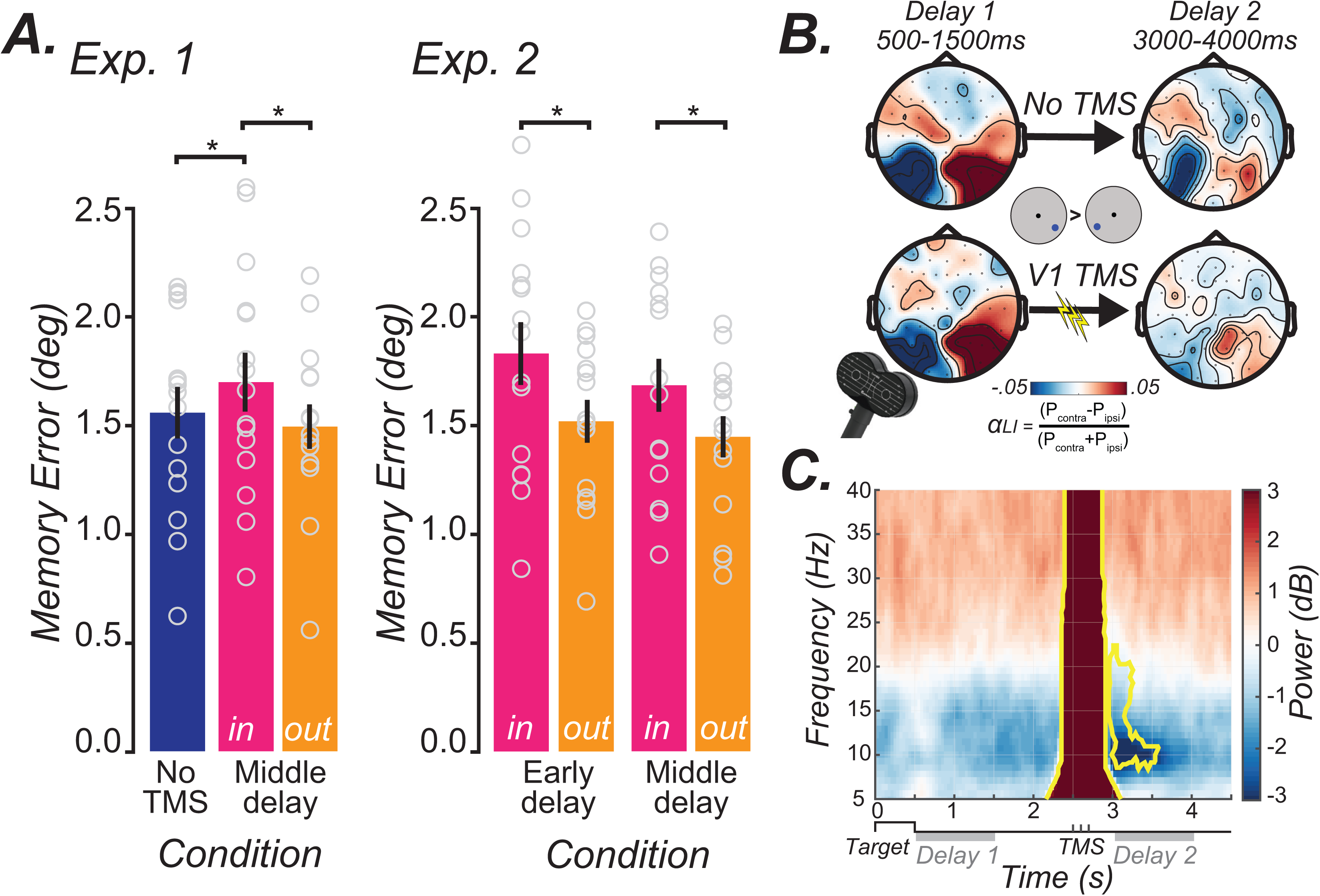
Effects of TMS to V1 on working memory. **A.** In Experiment 1, TMS applied to V1 during the middle of the delay period caused an increase in memory errors for targets in the phosphene field (in), compared to when no TMS was applied and to memory for targets in the opposite hemifield (out). In Experiment 2, we replicated the results from Experiment 1 by again showing that TMS to V1 during the middle of the delay caused a specific worsening of memory for targets in the phosphene field. Moreover, we found similar effects when TMS to V1 was delivered as soon as the visual target was extinguished (early) and when it was applied later in the middle of the delay (middle). Bars depict means, with SEM error bars, and circles are individual participants (n = 15). (* denote two-sided paired *t-*tests *p*<0.05). **B.** EEG collected during Experiment 1 were used to compute an alpha lateralization index (αLI), sensitive to the topography of alpha power in electrodes over occipital cortex related to the location of working memory targets. Without TMS, alpha topography tracked the remembered targets evidenced by a decrease in alpha power contralateral to the target (Delay 1). This pattern persisted late into the delay period (Delay 2). Following TMS applied to V1 during the middle of the memory delay (2000 ms), note how the strength of the lateralized alpha decreased (Delay 2). **C**. EEG time-frequency responses (TMS minus no TMS) during the working memory delay for occipital electrodes over the hemisphere being stimulated. Following the TMS evoked artifact (large red broadband increase), power significantly decreased in alpha and beta bands. Yellow outlines significant clusters of power differences between TMS minus no TMS (cluster-based permutation two-sided paired t-test, corrected for multiple comparisons, *p*<0.05). This further explains the effect of TMS on αLI. Gray lines along the timeline denote the Delay 1 and Delay 2 epochs in B. Source Data are provided in the Source Data file.

In Experiment 2, we aimed to both test the reproducibility of these results and how the timing of the stimulation impacts working memory performance. The same participants (n=15) returned and we again delivered TMS to V1 during the delay period of the memory-guided saccade task (Figure 1F). First, we perfectly replicated our findings from Experiment 1. TMS applied to V1 during the middle of the delay period caused greater memory errors for targets inside the phosphene field compared to targets in the mirrored hemifield (*t*(14)=2.742, *p*=0.007, *d*=1.47). Next, we asked how TMS applied during working memory maintenance compares to when applied during encoding (Figure 1F). TMS applied to V1 early in the delay (at the offset of the visual target) impaired memory accuracy for targets inside the phosphene field compared to targets outside it (*t*(14)=3.303, *p*=0.003, *d*=1.77). Although numerically larger, these errors caused by TMS at encoding were not statistically greater than those caused by TMS during maintenance (*t*(14)=1.248, *p*=0.095, *d*=0.67).

Next, since we used memory-guided saccades as our measure of working memory, we explored what aspects of the saccade metrics were affected by TMS. First, TMS applied during the middle of the delay (*t(14)*=-0.457, *p*=0.655, *d*=0.244) or during the early delay (*t(14)*=0.251, *p*=0.806, *d*=0.134) did not impact saccade reaction times (Supp Figure 2A). Similarly, TMS applied during the middle of the delay (*t(14)*=0.321, *p*=0.753, *d*=0.171) or during the early delay (*t(14)*=0.261, *p*=0.798, *d*=0.140) had no impact on the peak velocity of saccades (Supp Figure 2B). Therefore the effects of TMS we observed cannot be explained by a speed-accuracy tradeoff or slowing of saccades. Second, using regression we found that memory-guided saccade errors were driven by both radial (i.e., gain; No TMS *t(14)*=24.992, *p*=5.1487e-13, *d*=13.359; TMS outPF *t(14)*=19.789, *p*=1.2420e-11, *d*=10.577; TMS inPF *t(14)*=24.090, *p*=8.5177e-13, *d*=12.877) and tangential (i.e., angular; No TMS *t(14)*=7.822, *p*=1.7791e-06, *d*=4.181; TMS outPF *t(14)*=8.236, *p*=9.7491e-07, *d*=4.402; TMS inPF *t(14)*=7.214, *p*=4.4699e-06, *d*=3.856) error. Moreover, TMS condition did not impact the way in which saccade errors were decomposed into these components (radial - No TMS vs TMS *t(14)*=-1.732, *p*=0.105, *d*=0.926; radial - TMS inPF vs TMS outPF *t(14)*=0.651, *p*=0.525, *d*=0.348; tangential - No TMS vs TMS *t(14)*=0.214, *p*=0.834, *d*=0.114; tangential - TMS inPF vs TMS outPF *t(14)*=-0.4874, *p*=0.634, *d*=0.261). Therefore, both radial and tangential components were equally impacted by TMS. This is important because it suggests that targeting V1 with TMS caused errors that cannot merely be explained by some systematic motor factors (e.g., only hypometricity). Instead, they point to induced errors in memory that were highly specific to targets in the phosphene field.

Another possible explanation for our results is that the visual presence of phosphenes acted as distraction or somehow captured attention. Phosphenes were rarely reported by participants because of the brighter gray background during the experiment compared to the black used during phosphene mapping. When they were reported on rare occasions, the intensity of the TMS was reduced and participants no longer reported seeing phosphenes. Moreover, it is not clear what the prediction would be for such a visual distraction as behaviorally relevant, but not irrelevant, stimuli can impact working memory representations in early visual cortex ^21,22^. Nonetheless, we performed a control experiment in which participants were shown simulated phosphenes as three flickers matching the exact times of the three TMS pulses during the middle of the memory delay (See *Methods*). The pseudo phosphenes were low contrast noise patterns created to match subjective reports from our participants, and were shown at the same exact positions as the real phosphene fields experimentally defined (Supp Figure 4A). Memory errors were comparable for trials where pseudo phosphenes were presented on the same or opposite hemifield compared to the target (*t(9)*=0.671, *p*=0.5194, *d*=0.447) (Supp Figure 4B), therefore ruling out that our TMS effects were caused by rare distracting effects of visible phosphenes. Together, these results demonstrate that perturbations targeting V1 are reliable, retinotopically specific, and are not likely the cause of oculomotor disturbances or secondary effects of visual distraction.

We used electroencephalography (EEG) in Experiment 1 to measure the impact that TMS targeting V1 had on the characteristic asymmetry of alpha band topography over occipital cortex electrodes during working memory ^23^. Without TMS, alpha power decreased in the hemisphere contralateral to the memory target early in the delay, which we quantified with an alpha lateralization index (αLI) (Figure 2B, top). The lateralized alpha response persisted into the late portion of the delay, albeit lower in amplitude. On TMS trials, early in the delay the topography of alpha was indistinguishable from the no TMS trials before stimulation. Following TMS, the asymmetric alpha response was reduced compared to no TMS (Figure 2B, bottom). Next, we subtracted the EEG time-frequency responses of the no TMS trials from the TMS trials among the occipital electrodes over the hemisphere in which stimulation was applied (see *Methods*). Once the TMS artifact resolved (Supp Figure 3), we found significant reductions in alpha band power for about 500ms and beta band power for about 250ms (Figure 2C). Therefore, consistent with the increase in behavioral memory errors, TMS also disrupted a prominent EEG signature of working memory.

## Discussion

In summary, TMS targeting human V1 caused a deficit in working memory that was spatially restricted to the portion of the visual field perturbed by the electrical field induced by the TMS pulses. While previous studies testing how visual information is consolidated into working memory report that TMS to occipital cortex during the early encoding phase of working memory disrupts memory performance ^24–26^, we find that TMS during the maintenance phase produced deficits just as robust as those during encoding. The few other TMS studies of early visual cortex for working memory have been unable to confirm or refute whether it is necessary. For instance, TMS to occipital cortex only slowed working memory based decisions when applied after the delay period ^25^. In one study, TMS worsened working memory precision very early in the delay (100ms), but not later (400ms) or even during the presentation of the visual array ^26^. In another study, TMS had an effect in only 1 out of 6 conditions (high working memory load at 400ms into the delay) ^24^. In yet another study, 10Hz TMS actually improved working memory precision for gabor orientations ^27^. Finally, TMS to human MT disrupted working memory but only for the more prioritized of two memorized motion stimuli ^28^. Although we can only speculate about the various factors that might underlie the inconsistency of these results, we will highlight a few here. First, most of the previous studies were designed to mainly test the role of visual cortex in encoding or early consolidation, consistent with the notion that its role is limited to memory encoding and not maintenance. Second, methodological issues might have hampered clear interpretations. Here, we controlled the retinal specificity of the effects by placing stimuli in the phosphene fields during the early and middle of the memory delay. We applied strong triple pulses of TMS that produced robust effects. We used measures sensitive to small changes in working memory behavior. We also used EEG to confirm that our perturbations impacted neural indices associated with working memory. Bolstered by these experimental procedures, our findings provide direct causal evidence that early visual cortex may be necessary for working memory, beyond simply the visual encoding or consolidation of stimuli into memory.

As these results have important consequences for theories of working memory, we considered alternative explanations. For instance, perhaps higher-order areas known to be important for working memory, like the frontal and parietal cortices, were somehow affected by TMS targeting V1. We believe that this is unlikely. First, neither V1 or V2 have direct monosynaptic projections to the dorsolateral prefrontal cortex, area BA46 in the macaque ^29^. The vast majority of connections between V1 and lateral intraparietal area (LIP) or frontal eye field (FEF) are not monosynaptic ^29^. For instance, monkey V1 is mainly connected to LIP indirectly through several synapses ^30^. Second, PET and fMRI studies of concurrent TMS suggest that distal areas affected are only those that have monosynaptic connections to the brain target of the TMS ^31,32^. Third, electrical microstimulation of macaque FEF without concurrent visual stimulation does not evoke BOLD activity in V1 ^33^. Fourth, TMS perturbs spiking activity only within a 2mm radius, and synaptic potentials only within a slightly larger extent in the macaque cortex ^34^. This evidence together suggests that the effects we observed here are due to highly localized effects of TMS to V1 and perhaps V2d.

We must also acknowledge two caveats to our conclusion that V1 is necessary for working memory. One, while we targeted retinotopically defined V1, perhaps the effects we observed were due to the propagation of electrical activity to other distant brain areas like the prefrontal cortex. While by far the strongest electrical effects are localized to brain areas directly targeted by TMS, one would assume that smaller effects might propagate to areas a single synapse away. Based on the macaque brain, neither V1 or V2 has direct monosynaptic connections with the dorsolateral prefrontal cortex, and the overwhelming majority of connections between V1 to LIP and FEF are not monosynaptic ^29,30^. Therefore, if our results were due to indirect effects of TMS on areas monosynaptically connected to V1, they are most likely limited to areas extensively connected with V1, such as V2, V3, and V4 and subcortical areas such as the lateral geniculate nucleus. Indeed, no evidence exists that the indirect effects of TMS on distant brain areas have effects on behavior. Rather, the effects on neuronal spiking caused by TMS are extremely focal (∼2mm) ^34^ and single finger contractions can be individuated by TMS displacements of only ∼1mm across cortex ^35^. Two, TMS targeting V1 worsened the accuracy of memory-guided saccades, which we interpreted as an impact on visuospatial working memory. However, other possible cognitive or motor behaviors may have been affected, and thus masqueraded as a memory deficit. Although spatial working memory and spatial attention are thought to rely on overlapping neural mechanisms ^36,37^, TMS to V1 surprisingly does not affect endogenous covert attention ^38^. Additionally, perhaps TMS targeting V1 somehow impacted the oculomotor metrics used to plan saccades ^39,40^. We do not believe that the results are consistent with this possibility as the memory-guided saccade errors induced by TMS were highly specific to memory targets inside the contralateral phosphene field. Saccade response times and peak velocity were not affected. Moreover, the types of memory-guided saccade errors were not systematic; TMS impacted both the gain and angle of saccade errors (again, only in the contralateral visual field). Nonetheless, future research should extend our results by testing if more complicated forms of visual working memory depend on V1, taking into account the difficult challenges in establishing causality ^41^.

Our findings provide compelling support to human neuroimaging studies that consistently report that the visual properties of memoranda stored in working memory can be decoded from the patterns of activity in V1 during retention intervals ^7,9^. Moreover, they align with neurophysiological recordings from macaque V1 demonstrating stimulus selective activity during working memory delays ^42–45^. They provide a key piece of evidence that had been missing, but needed to directly test the sensory recruitment hypothesis of working memory. Indeed, we demonstrate that the neural mechanisms housed in V1 (and perhaps V2d) are necessary to maintain accurate working memory representations. Therefore, the question is no longer if visual cortex is necessary for working memory, but now it is by what mechanisms does it contribute to working memory? Assuming indirect effects do not account for our results, they are consistent with models in which the neural mechanisms housed in V1 (and perhaps V2d) play an important role in maintaining accurate working memory representations. Given the accumulation of evidence from neuroimaging, electrophysiology, and TMS, we argue that the question should no longer be about if visual cortex plays an important role in working memory, but instead should be about by what mechanisms does it contribute to working memory?

## Methods

### Participants

All experiments were conducted following the institutional guidelines of New York University and the regulations of the institutional review board (IRB).Twenty-seven neurologically healthy humans (12 female, 15 male; mean age: 28, range: 21-55) were recruited to participate in multi-session TMS studies. All participants had normal to corrected normal vision and were screened for TMS eligibility and excluded from the study if they had any brain-related medical issues or were currently taking certain drugs (e.g. antidepressants, amphetamines, etc.). All participants provided written, informed consent and were compensated $50 for each session, regardless of whether it involved TMS (since the order of TMS was randomized across participants). Three participants found the procedures uncomfortable and withdrew. Five participants were withdrawn because we could not measure their gaze well enough (e.g., poor pupil lock, excessive blinking, lid closures, and dry/watery eyes). Since gaze was our main dependent variable, we could not study these participants. One participant was excluded because they did not follow instructions and made saccades only after the visual feedback was given, and thus not based on memory. Three participants were excluded because we could not reliably evoke phosphenes in the contralateral hemifield. The results, therefore, were based on fifteen participants (7 female, 8 male; mean age: 31, range: 21-55) who completed the full study. Two of the participants were authors and the rest were naive to the purpose of the study.

### Rapid Serial Visual Presentation (RSVP) Task and Retinotopic Mapping

Anatomical and functional brain scans for pRF mapping were collected at New York University Center for Brain Imaging (CBI) using a 3T Siemens Prisma MRI scanner using a Siemens 64-channel head/neck radiofrequency coil. Volumes were acquired using a T2*-sensitive echo planar imaging pulse sequence (repetition time (TR), 1300 ms; echo time (TE), 36 ms; flip angle, 66°; 56 slices; 2 mm x 2mm x 2mm voxels). High-resolution T1-weighted images (0.8 mm x 0.8 mm x 0.8 mm voxels) were collected at the end of the session, with the same slice prescriptions as for the functional data, and used for registration, segmentation, and display. Multiple distortion scans (TR 6,690 ms; TE 63.4 ms; flip angle, 90°; 56 slices; 2 mm x 2 mm x 2 mm voxels) were collected during each scanning session.

To define visual field maps, each participant underwent retinotopic mapping in the MRI scanner, following previously established procedures ^19^. Participants performed a RSVP task composed of sweeping bars of stimuli that swept across the screen in swift and discrete steps, moving in four possible directions: leftward to rightward, top to bottom, rightward to leftward, or bottom to top. Each bar contained six images of colorful objects. At the start of each block, participants were shown the target object and were instructed to make a button press every time the target object was present anywhere within the array while maintaining fixation. The participants received online feedback with the fixation cross turning green indicating their response was accurate. The inter-stimulus interval was 400 ms and was titrated at the end of each sweep to match participants’ accuracy to 70-80%. The data were monitored online for fixation breaks and blocks where participants broke fixation were eliminated from pRF model-fitting. Participants performed around 10 runs of the task, with 8 bar sweeps per run.

Resulting BOLD time series were fitted with a population receptive field (pRF) model with compressive spatial summation ^46,47^, and visual field maps, including V1, were drawn manually using afni and suma. Specifically, we visualized the polar angle and the eccentricity maps on the cortical surface, thresholded to include only voxels for which the pRF model explained more than 10% of the variance. These ROIs were then inflated using volumetric data as grid parent and used for targeting with TMS (as described below).

### Transcranial Magnetic Stimulation (TMS)

The T1-weighted images of participants were used to reconstruct brain and skin surfaces to localize targets for TMS using a frameless stereotaxic neuronavigation system (Brainsight, Rogue Research Inc., CAN) combined with Polaris Spectra (Northern Digital, USA) infrared cameras that tracked markers positioned on the TMS coil and on the participant’s head. TMS was applied using a MagVenture MagPro X100 stimulator with a MCF-B70 static cooled coil. TMS pulses were triggered and coupled with EEG using custom MATLAB and Python scripts (https://github.com/clayspacelab/EEG_TMS_triggers). We identified V1 in each participant from fMRI-based retinotopy (see *Retinotopic Mapping* above) and overlaid the target on the brain surface in Brainsight. We initially randomized whether the participant would receive TMS to the left or right V1. However, if we had difficulties evoking reliable phosphenes we switched to the other hemisphere. Once we identified a putative target that evoked phosphenes, we then determined the phosphene threshold and mapped the spatial extent of the phosphene (see *Phosphene Mapping* next).

### Phosphene Mapping

Participants were first screened for their ability to detect phosphenes in a dimly lit room by placing the TMS coil over retinotopically defined V1 and applying 7 pulses at 20 Hz. The coil was oriented with the handle away from the interhemispheric fissure to induce currents along the caudal-rostral axis. We made small changes to the position and angle of the TMS coil until we identified a target location that reliably induced phosphenes. If phosphenes were inconsistent or weak, we tried stimulating V1 in the other hemisphere. Next, we then measured the spatial extent and borders of phosphenes. Participants used a computer mouse to encircle the perceived phosphene on several trials until consistency was achieved, defined as at least three trials with overlapping phosphenes. For each participant, we defined their phosphene field (PF), which we derived from the overlapping area of the phosphene drawings. The PF was the area of visual space consistently impacted by TMS to V1 at the given coil position. The area of the PF was determined by the following criteria. First, the centroid was calculated as the center of the overlapping area. Second, the angular extent spanned the full distance along an arc running through the centroid. Third, the radial extent was defined as a 2 degree band centered at the centroid. In the experiment, we placed memory targets inside the area of the defined PF (see *Memory-Guided Saccade Task* below). Once a reliable PF was identified, we estimated their phosphene threshold of TMS intensity. Starting at 30% of maximum stimulator output we delivered 7 pulses at 20 Hz and recorded if a phosphene was evoked. We increased the stimulation intensity by 10% until phosphenes were reliably evoked, then reduced stimulation intensity by 5% to find the threshold defined as reported phosphene in 5 out of 10 trials. In the main experiment, we used a stimulation intensity at 110% of that threshold, but only applied 3 pulses at 20 Hz. Note that the phosphene thresholds were determined using 7 pulses and in a dark room looking at a screen with a black background. During the main experiment, participants received 3 pulses looking at a brighter screen with gray background. Therefore, the phosphenes were rarely perceived. On rare occasions when participants did report phosphenes, usually noted during the first couple trials of the first block, the intensity was lowered until participants no longer perceived phosphenes. See below for details of Control Experiment 3, which was designed to test if the visual perception of phosphenes impact memory.

### Memory-Guided Saccade Task

#### Experiment 1: TMS during working memory maintenance. Middle Delay

Participants performed a memory-guided saccade (MGS) task. Each block consisted of 40 trials and began with a fixation cross, followed by a target presented for 500ms inside the PF, or the mirrored hemifield of the PF, randomized across trials. Participants maintained in working memory the location of the target over a 4s retention interval while fixating a central circular fixation stimulus. When the fixation stimulus turned green, they made a saccade, guided by their memory alone, to the location of the target. After 850ms, the target reappeared in green serving as feedback. Experiments 1 and 2 were conducted over four days to minimize both fatigue and the number of TMS pulses participants experienced in a given day. On day 0, participants performed a 1 hour TMS session of phosphene thresholding, phosphene mapping (described above), and practice of MGS task. On days 1-3, participants performed 5 blocks (200 trials; 600 TMS pulses) of the MGS task during which their brain activity was recorded using EEG (see *Electroencephalography* below). On 2 of these 3 days, TMS was applied to V1 during the middle (at 2s) of the 4s long delay (what we refer to as *Middle Delay)*. On the other day, no TMS was applied. For each participant, we randomized the order of the days on which TMS was applied.

#### Experiment 2: TMS during working memory encoding. Early and Middle Delay

Each participant returned on another day to complete two sessions of the MGS task, while TMS was applied to the V1 target area previously identified. EEG was not collected. The experiment was identical to that described in Experiment 1, with the only difference being that TMS was applied either at target offset (what we refer to as *Early Delay*) or in the middle of the delay (what we refer to as *Middle Delay*). The order of sessions was randomized across participants. Each session lasted 40 minutes and participants performed 5-7 blocks of MGS task (described above), followed by a mandatory 10-minute break before they performed the next session lasting 40 minutes.

#### Control Experiment 3: Simulated phosphenes during working memory delay

Ten participants returned for another day of study to complete one session of MGS task where TMS during the middle of the delay was replaced by simulated phosphenes. Specifically, the visual appearance of phosphenes was simulated in the phosphene field reported by the participants during the phosphene mapping task. A white noise mask was added to the phosphene field area, presented as three brief flickers during the middle of the delay, simulating the three pulses of TMS during experiments 1 and 2. In order to control for any potential hemispheric differences in performance, simulated phosphenes were presented either on the same or opposite hemifield compared to the target. The purpose of this experiment was to test the degree to which perceived phosphenes could account for memory errors. Each participant completed 10 blocks of MGS in one session that lasted 75 minutes and participants were compensated $15 for their time.

### Eye-tracking and Visual Display

Participants were seated in a dim-lit room with the head positioned on a chin-rest 56 cm away from a gamma-calibrated ViewPix monitor (120Hz refresh rate, 1920*1080 resolution) controlled by an Ubuntu video card. Stimuli were presented using MATLAB and Psychtoolbox ^48^. An Eyelink 1000 Desktop Mount (SR Research) was used to record the right eye position at 1000 Hz. For all participants, a 9-point calibration was performed at the start of the session and repeated whenever the participant took a break by moving the head from the chin-rest. The data were then preprocessed using iEye (https://github.com/clayspacelab/iEye), an in-house gaze-analysis toolbox written in MATLAB. The preprocessing involved multiple steps. First, blinks were identified as pupil diameter lower than 1 percentile. Gaze data 100 ms before and after blinks were removed. Next, a Gaussian smoothing kernel was applied to gaze traces to get an estimation of velocity. This was used to detect saccades, which were defined as gaze transitions with velocities greater than 30 deg/s, having an amplitude greater than 0.25 degree, and a duration of at least 7.5 ms. The data were then corrected for any instrumental drift by demeaning the gaze traces during fixation and delay epochs. Lastly, we also performed calibration using the corrective saccades during the feedback for any instrumental or head-movement-related drift that might have occurred in a trial. Given eye-tracking was the sole metric of behavior, these extra precautions ensured quality gaze data. Trials where stimulus timings were off (∼0.2%) participants made large errors (>10dva) (∼4.6%), where the saccade reaction time was less than 150ms (likely suggesting participants started saccade during delay epoch) (∼0.54%), and trials where participants blinked during the saccade or the eye tracker lost saccade traces during experimental setup failures (∼8.9%) were eliminated from analysis.

### Electroencephalography (EEG)

We used a BrainProducts 64-channel actiChamp system with low-profile active electrodes for recording EEG activity. Data were sampled at 1000Hz, with the ground placed at the center of forehead above nasion and Cz was used as the online reference. EEG data were analyzed using Fieldtrip toolbox ^49^. Any trials with errors in timing of the TMS pulse were detected (less than 2 out of 400 trials on average per session) and removed from analysis. Then, the data for each session were band-pass filtered between 0.5-50 Hz using a two-pass butterworth filter of filter order 4. The data were then screened for bad channels. Bad channels were identified as ones that exhibited more than 90 percentile of median variance amongst all channels, were flat (std < 0.01 µV), or were extremely noisy (std > 100 µV) ^50^. The cleaned data were then offline re-referenced to the common average. Next, owing to long delays in our working memory task, the data were heavily contaminated with blink artifacts. Independent Component Analysis (ICA) was performed using the ‘fastica’ algorithm in Fieldtrip and components exhibiting muscle twitches, eye-movement artifacts and saccades were carefully examined and eliminated. Bad channels removed earlier were then interpolated using a weighted interpolation of neighboring electrodes. The data were then epoched and downsampled to 200Hz. The epoched data were then examined for bad trials which were then eliminated (less than 0.5% on average). Additionally, time-frequency analysis (TFA) was conducted to investigate spectral dynamics. The analysis was performed using the wavelet method. The frequency range was set from 2 to 40 Hz, discretized into 53 linearly spaced frequency bins. The number of cycles for the wavelet analysis varied linearly with frequency, ranging from 4 to 15 cycles across the specified frequency range. Subsequently, the power values were converted to decibel (dB) scale. Alpha Lateralization Index (αLI) was computed as the ratio of power difference between the contralateral and ipsilateral activity for each trial from the occipito-parietal electrodes (O1, O2, PO3, PO4, PO7, PO8, P1, P2, P3, P4, P5, P6, P7, P8) to the total power across all occipito-parietal electrodes. αLI = (P_contra -_ P_ipsi)_ / (P_contra +_ P_ipsi)_ where, P_contra w_as power in an electrode on trials where stimulus was present in contralateral hemifield, and P_ipsi w_as power in an electrode on trials where stimulus was present in ipsilateral hemifield.

### Electrical field simulations

Simulation of Noninvasive Brain Stimulation (SimNIBS) ^20^ 4.0 was used to estimate the electric field induced in the brain based on anatomical scans and the target location chosen for each participant as described above. Using CHARM-based head-segmentation ^51^ and target locations, E-field estimates were computed for each individual brain taking into account conductivities of different tissue types. The model uses the coil specifications provided by MagVenture to estimate the magnetic field induced by the coil. For each participant, the exact placement of the TMS coil, including the rotational angle was input to the model. This allowed us to create biophysical models of the amplitude and extent of the electric field induced by TMS to V1 for each participant.

### Statistical Analyses

Data analysis and statistics were performed using Python 3.11.5, Numpy 1.26.2, Scipy 1.11.4^52^ for behavioral statistics and Fieldtrip to perform cluster-based statistical analyses for EEG. No statistical methods were used to predetermine sample size but we used sample size estimates from similar previous studies^38^. To ensure a reliable measure of behavior, participants and trials were eliminated as described in “Participants” and “Eye-tracking and Visual Display” sections. Participants were carefully screened to be eligible for the study and hence the investigators were not blinded to the selection of the participants. However, the assignment of perturbation was randomized across participants and the investigators were blinded prior to the onset of the session.

For behavioral data, we randomly shuffled the trial labels for TMS conditions (No TMS, Early TMS, or Middle TMS) and target location (inside PF or outside PF). For each permutation, we computed the group average for each condition, thereby generating a null distribution of *t*-statistics. The observed *t*-statistic was then compared against this null distribution to obtain the *p*-value. In order to assess the contribution of radial and tangential components to saccade errors, we first *z*-scored radial, tangential and absolute errors and performed an ordinary least squares regression using statsmodel OLS to estimate the contribution of each component to the total error, separately for each subject and condition. We then performed repeated-measures ANOVA using statsmodel AnovaRM to assess the differences in the contribution of components to total error across conditions.

For time-frequency EEG analyses, we employed a cluster-based permutation approach to perform paired *t*-tests at each time-frequency bin using *ft_freqstatistics* function in Fieldtrip. The primary alpha level was set to 0.05, while the cluster threshold was set at cluster-alpha of 0.1. Significance of the clusters was assessed using Monte Carlo method with 1000 random permutations.

### Data Availability

Source data to reproduce bar plots in all figures are provided as a Source Data file. All data used in this study is made publicly available at https://osf.io/hf5a4/. The published dataset contains preprocessed and de-identified behavioral data across both experiments, preprocessed time-frequency analyzed EEG data, and the TMS-efield simulations and ROIs for each participant. Any raw data can be made available upon request.

### Code Availability

Code used to trigger the TMS and EEG systems is available at https://github.com/clayspacelab/EEG_TMS_triggers. The toolbox iEye used to process eye-tracking data is available at https://github.com/clayspacelab/iEye. And the codes required to replicate figures from data are made available at https://github.com/clayspacelab/Dake_Curtis_TMSV1_2025.

## Acknowledgments

We thank the staff of the Center for Brain Imaging (CBI) at New York University (NYU), and Yuyang Xu and Qingqing Yang for help with data collection. We are also grateful to Nathan Tardiff and Zhengang Lu for their comments on drafts of the manuscript. Funded by NIH R01 EY-016407 and R01 EY-033925 to C.E.C.

## Author Contributions

M.D. performed the experimental studies, analyzed the data, and wrote the manuscript. C.E.C. conceived the experiments, supervised the work, and wrote the manuscript.

## Competing Interests

The authors declare no competing interests.

## Notes

https://osf.io/hf5a4/

### Competing Interest Statement

The authors have declared no competing interest.

### Summary of Updates

Formatting changes and responses to reviewers.

