## Supplementary Figure 1, 2, 3, 4 for "Perturbing human V1 degrades the fidelity of visual working memory"

### Supplementary Materials

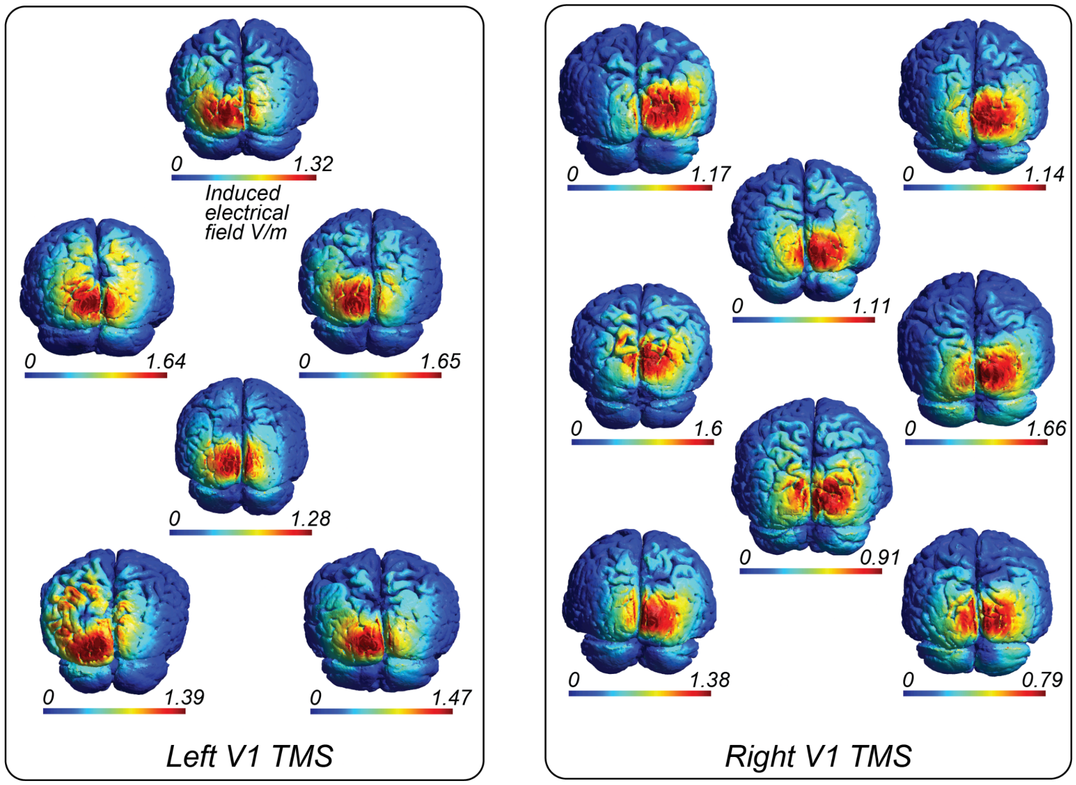


***Supplementary Figure 1 — Induced electric field simulations for individual participants:*** *Induced E-field (V/m) in individual brains as predicted by SimNIBS on native brains for a single pulse of TMS applied at the target location used for each participant. (Left) For participants with left V1 stimulated. (Right) For participants with right V1 stimulated.*

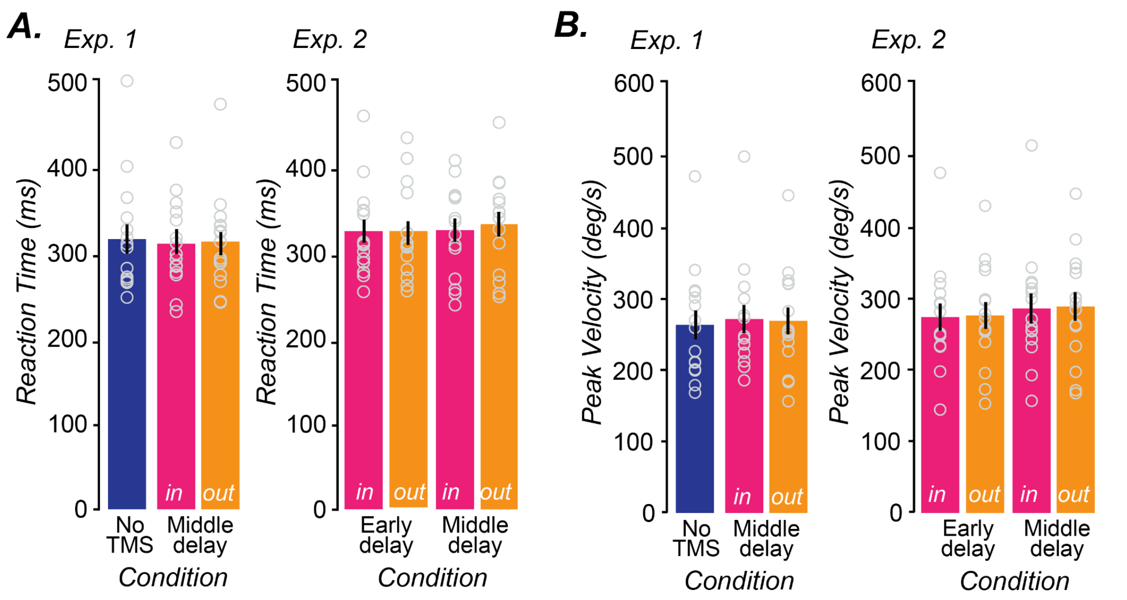


***Supplementary Figure 2 — Effect of TMS on Saccade RT and Peak Velocity*** *A) Saccade reaction times (sRT) (in ms) for different conditions. There was no impact of TMS on sRT either when TMS was applied during early or middle delay relative to No TMS (paired t-test: p > 0.05; n=15 humans). Similarly, there was no difference in sRT for when targets were present inside or outside PF (paired t-test: p > 0.05; n=15 humans) when TMS was applied during either the early or middle part of the delay. B) Saccade peak velocity (in deg/s) for different conditions. There was no impact of TMS on peak velocity either when TMS was applied during early or middle delay relative to No TMS (paired t-test: p > 0.05; n=15 humans). Similarly, there was no difference in peak velocity for when targets were present inside or outside PF (paired t-test: p > 0.05; n=15 humans) when TMS was applied during either the early or middle part of the delay. Data provided in the Source Data file.*

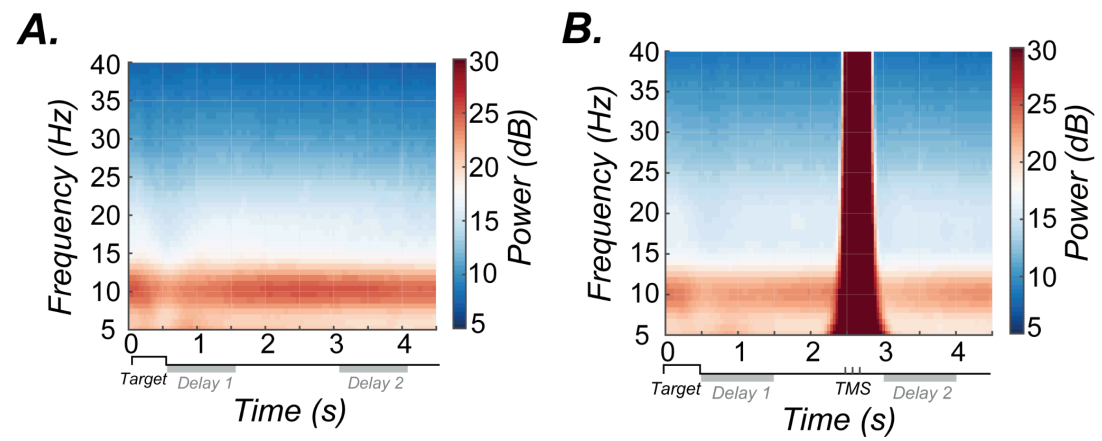


***Supplementary Figure 3 — EEG Time-Frequency Responses during Delay Period*** *A) No TMS, B) TMS, shows the time-frequency responses averaged across participants (n=15 humans) for occipital electrodes under the TMS site. Note that TMS induces a broad-band frequency change beginning with the onset of the TMS pulse and it lasts for 500ms after the TMS pulse ends, after which there is a brief dip in alpha frequency band, characterized strongly by the change in αLI depicted in Figure 2.*

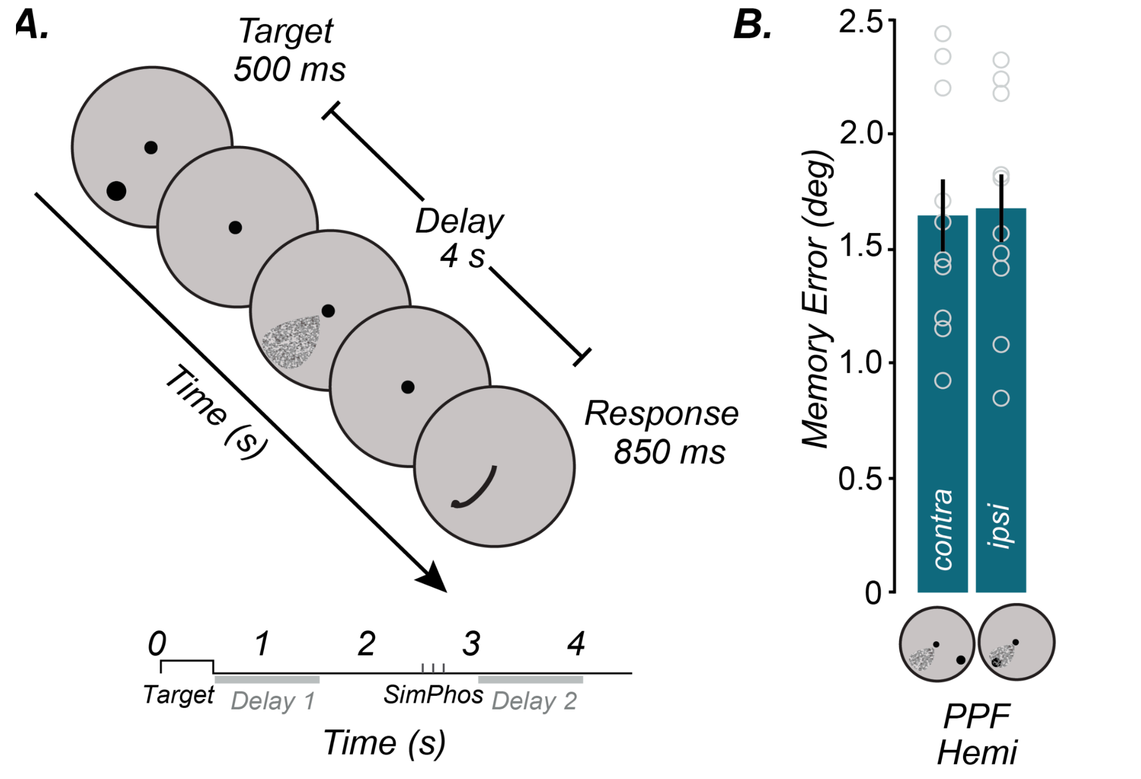


***Supplementary Figure 4 — Effect of simulated phosphenes on MGS task performance*** *A) Schematic of simulated phosphenes during MGS task. Note that the pseudo phosphene was presented at a lower contrast similar to that reported by participants. The placement and area was exactly matched to each participant's experimentally derived phosphene fields (referred to as pseudo-phosphene fields (PPF)). B) There was no impact of simulated phosphenes on memory errors either when targets were presented inside PPF (ipsi) or outside PPF (contra)) (paired t-test: $t(9) = 0.6705$, $p = 0.5194 > 0.05$, $d = 0.45$; $n = 10$ humans). Data provided in the Source Data file.*

***Supplementary Table 1*** **— Participant demographics and stimulation parameters**

| Participant | Handedness | Hemisphere stimulated | Phosphene Threshold (% MSO) | Centroid of Phosphene [eccentricity (dva), polar angle (degrees)] |
|---|---|---|---:|---|
| 01 | Right | Left | 40 | (14.2, 352.0) |
| 02 | Right | Right | 42 | (16.8, 192.9) |
| 03 | Right | Left | 51 | (11.6, 337.4) |
| 04 | Right | Right | 62 | (13.0, 193.8) |
| 05 | Left | Left | 52 | (14.3, 336.1) |
| 06 | Right | Right | 42 | (6.2, 186.8) |
| 07 | Right | Left | 50 | (7.48, 341.9) |
| 08 | Right | Right | 40 | (13.8, 208.6) |
| 09 | Left | Right | 37 | (16.0, 196.1) |
| 10 | Right | Left | 44 | (12.2, 334.4) |
| 11 | Right | Right | 43 | (11.5, 183.6) |
| 12 | Right | Right | 48 | (14.3, 200.1) |
| 13 | Right | Right | 43 | (16.2, 215.8) |
| 14 | Right | Left | 35 | (6.6, 345.7) |
| 15 | Right | Left | 43 | (12.8, 330.0) |
